# A Comprehensive Model to Differentiate Spontaneous, Drug-induced, and CSCs-related Drug Resistance

**DOI:** 10.1101/2024.05.04.592529

**Authors:** Kaixin Zheng

## Abstract

Drug resistance is a pivotal research area in oncology research, yet the integration of multiple sources of resistance into the evolution of drug resistance remains elusive. This study investigates dynamics of drug resistance in chemotherapy utilizing a mathematical model given a treatment protocol. The model categorizes drug resistance into spontaneous, drug-induced, and cancer stem cells (CSCs)-related types. Introducing a novel mathematical framework, this study incorporates explicit dosage-dependent terms to design tailored treatment strategies. A comparative analysis contrasts continuous constant therapy with periodic bolus injection. Virtual patients’ survival times are assessed under baseline dosages for both therapies, revealing the interplay between constant dosage in continuous therapy and maximum dosage in bolus injection on survival time. Our findings demonstrate that, at equivalent cumulative dosages, bolus injection markedly extends patient survival. Furthermore, a potentially bimodal relationship emerges between bolus injection efficacy and maximum dosage, suggesting that two optimal bolus injection strategies may hold.

## Introduction

Chemotherapy stands a primary treatment method for cancer [1]. However, its effectiveness is limited by tumor resistance to drug: 90% of chemotherapy failure is related to drug resistance [2]. Microenvironmental and molecular factors play a major role in most drug resistance development [2, 3]. For instance, hypoxia, as a microenvironmental factor, is crucial in the efficacy of certain anticancer drugs and radiation therapy [4, 5]. An example of a molecular factor is the over-expression of the MDR1 gene, which contributes to the phenomenon of multidrug resistance (MDR) [6].

In cancer treatment, drug resistance within a particular phenotype of cancer cells can be spontaneous, drug-induced, or a combination of both [1]. Spontaneous drug resistance occurs if a portion of the cancer cell population is genetically drug-resistant, independent of drug administration. Drug-induced resistance involves mutations induced by drug in certain cancer cells, rendering them resistant. Differentiating spontaneous and drug-induced resistance is extremely challenging. In 1997, Leith et al. identified the crucial role of MDR1 overexpression in causing older acute myeloid leukemia patients to respond poorly to chemotherapy compared to younger patients [7]. However, until 2013, this resistance was eventually determined to be drug-induced: Pisco et al. demonstrated through experimental and mathematical verification that in HL60 leukemic cells treated with vincristine, resistance is predominantly drug-induced rather than a result of the selection of pre-existing MDR1-expressing cells [8].

Distinguishing drug resistance is further complicated by the existence of cancer stem cells (CSCs). These cells are recognized for their significance in not only cancer initiation, but also in the cancer cell population maintenance. CSCs are know to be less susceptible to drug treatment [9]. The fraction of the CSCs in the tumor negatively regulates the cancer treatment response [10]. Many current models [11–16] for differentiating drug resistance focus solely on sensitive and resistant differentiated cancer cells. Given the prevalence of CSCs in numerous cancers [17], it’s essential to include them in understanding drug resistance development and designing effective treatment strategies.

Bladder cancer, the fourth most common malignancy in men and also prevalent among women, is strongly associated with factors such as advanced cigarette smoking, male sex, and aging [18]. In the context of bladder cancer chemotherapy, understanding the evolution of tumor resistance involves integration of spontaneous, drug-induced, and CSCs-related resistance in understanding tumor resistance evolution. Drug-sensitive bladder cancer cells have been found to spontaneously transit to cisplatin-resistant phenotype [19]. Cho et al. demonstrated treatment-induced multidrug resistance (MDR) in bladder cancer [20]. In addition to the well-known enrichment of CSCs in bladder cancer chemotherapy [21], Kurtova et al. revealed that in vivo, bladder cancer CSCs contribute to therapeutic resistance by actively proliferating to repopulate residual tumors between chemotherapy cycles [22].

In this study, we want to quantitatively model three key aspects: 1. enrichment of CSCs after chemotherapy; 2. differentiating spontaneous, drug-induced, and CSCs-related drug resistance; 3. efficacy of dosage-dependent treatment.

## Method

### Model Formulation

To explain CSCs enrichment and CSCs-related resistance in bladder cancer, Weiss et al. [23] proposed an ODE model (Equation 1) of drug-sensitive CSCs, *C*_*S*_, drug-resistant CSCs, *C*_*R*_, drug-sensitive differentiated cancer cells (DCCs), *D*_*S*_, and drug-resistant DCCs, *D*_*R*_:

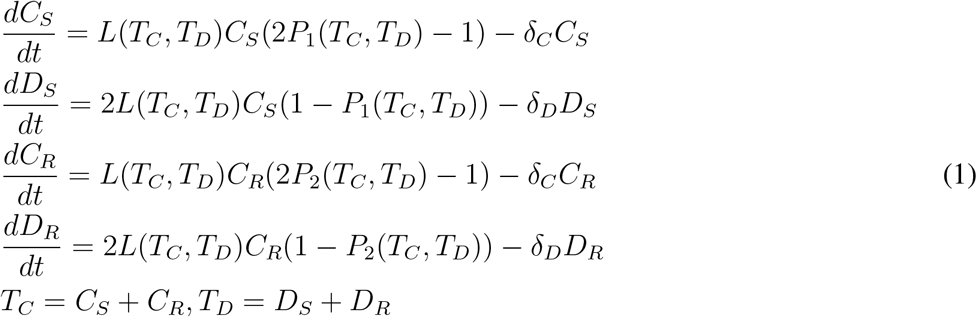

In Equation 1, proliferation rate *L*(*T*_*C*_, *T*_*D*_) is assumed to be function of population of CSCs, *T*_*C*_, and population of DCCs, *T*_*D*_. Regardless of phenotype, CSCs and DCCs have identical death rate, *δ*_*C*_, *δ*_*D*_ respectively. CSCs self-renewal probability in wild-type, *P*_1_(*T*_*C*_, *T*_*D*_) is assumed to be higher compared with that in mutated type (resistant type), *P*_2_(*T*_*C*_, *T*_*D*_).

Weiss model does not have an explicit drug-dosage-dependent term and the form of proliferation rate is undetermined. To address these issues, we adopt a similar approach to Greene’s framework model [15] to differentiate spontaneous and induced resistance: 1. Log-kill hypothesis [24] suggests that partial death rate caused by the drug is proportional to both the effective dosage and the cell population; 2. In the absence of treatment, the tumor is assumed to grow logistically; 3. There exists irreversible transition from sensitive phenotype to resistant phenotype. The last transition consists of two parts: first part is spontaneous and only proportional to sensitive population. The second part is drug-induced which is proportional to both sensitive population and effective dosage.

To further simplify our model, we assume self-renewal probability of CSCs are independent of CSCs and DCCs populations. Eventually, we propose a new model:

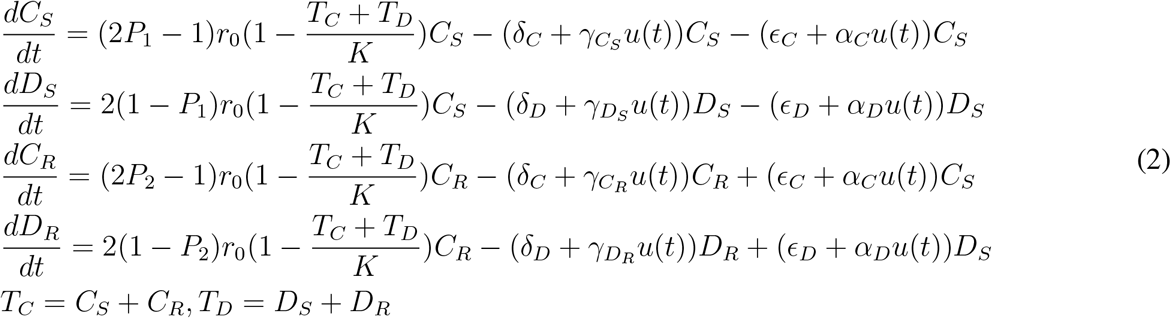

*u*(*t*) is the non-negative effective drug dosage applied at time t. *r*_0_ is growth rate for logistic growth. *K* is carrying capacity for the tumor. For subpopulation *X* = *C*_*S*_, *D*_*S*_, *C*_*R*_, *D*_*R*_, term *γ*_*X*_*u*(*t*)*X* represents the rate of drug-induced death, with cytotoxicity constant *γ*_*X*_ respectively. (*ϵ*_*Y*_ + *α*_*Y*_ *u*(*t*))*Y*_*S*_ is the transition rate between *Y*_*S*_, *Y*_*R*_ where *Y* = *C, D. ϵ*_*Y*_ *C*_*S*_ represents spontaneous drug resistance inside sub-population. *α*_*Y*_ *u*(*t*)*Y*_*S*_ is the drug-induced resistance with induction rate *α*_*Y*_. All parameters are non-negative.

### Non-dimensionalization and Modification

We rescale four sub-populations (joint) carrying capacity K, and time t by cell growth rate *r*_0_:

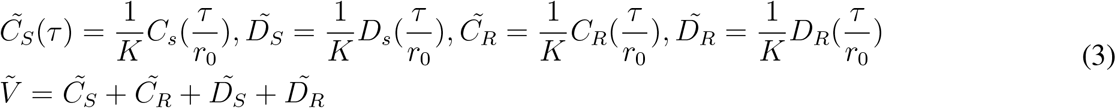

After rescaling all parameters by 1*/r*_0_, Equation 2 can be rewritten:

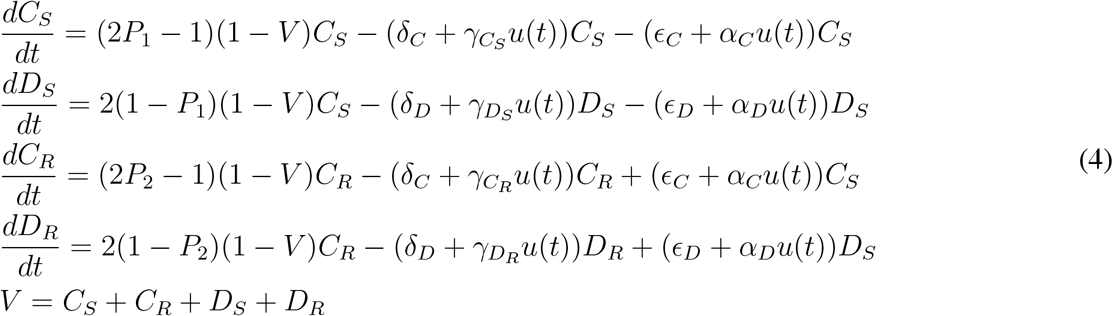

Biologically, if we assume volume of tumor is proportional to cancer cell population, *V* is the dimensionless volume of the tumor.

### Treatment Protocol

In this study, all simulations are conducted with dimensionless parameters and dimensionless initial conditions under a virtual treatment *u*(*t*). At *t* = 0, tumor volume *V* = *V*_*d*_ which is a detectable tumor volume. At *t* = *t*_*C*_, which is called survival time, tumor volume reaches *V* = *V*_*C*_, which is lethal for patients, and the treatment ends. The efficacy of treatment is quantified by duration of treatment, critical time, *t*_*C*_. The proportion of CSCs, both sensitive and resistant, relative to the tumor volume, V, is evaluated by proportion function:

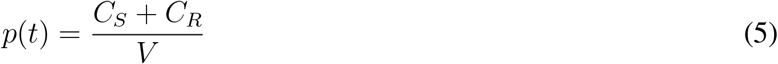

### Parameter Constraints and Baseline Parameters

If we choose *u*(*t*) = *O*(1), because spontaneous mutation rate is known to be significantly small [25, 26]. We may assume *ϵ*_*Z*_ ≪ *α*_*Z*_, *Z* = *C, D*. We can also reasonably assume *δ*_*Z*_ ≪ *γ*_*Z*_, *Z* = *C, D* because sensitive phenotype should be mainly killed due to drug administration or chemotherapy becomes unnecessary. Compared with DCCs, CSCs exhibit enhanced DNA repair capabilities through multiple mechanisms such as efficient engagement of the DNA damage response (DDR) pathway, activation of cell cycle checkpoints, and prolonged residence in the quiescent G0 phase of the cell cycle [27]. Therefore, we assume *δ*_*C*_ *< δ*_*D*_, *ϵ*_*C*_ *< ϵ*_*D*_, *α*_*C*_ *< α*_*D*_. Recalling CSCs are less susceptible to chemotherapy and expectations for resistant phenotype, we assume 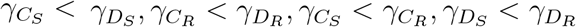.

In our Results section, unless specified, initial condition and all parameters are as listed below in Table 1.

**Table 1:**
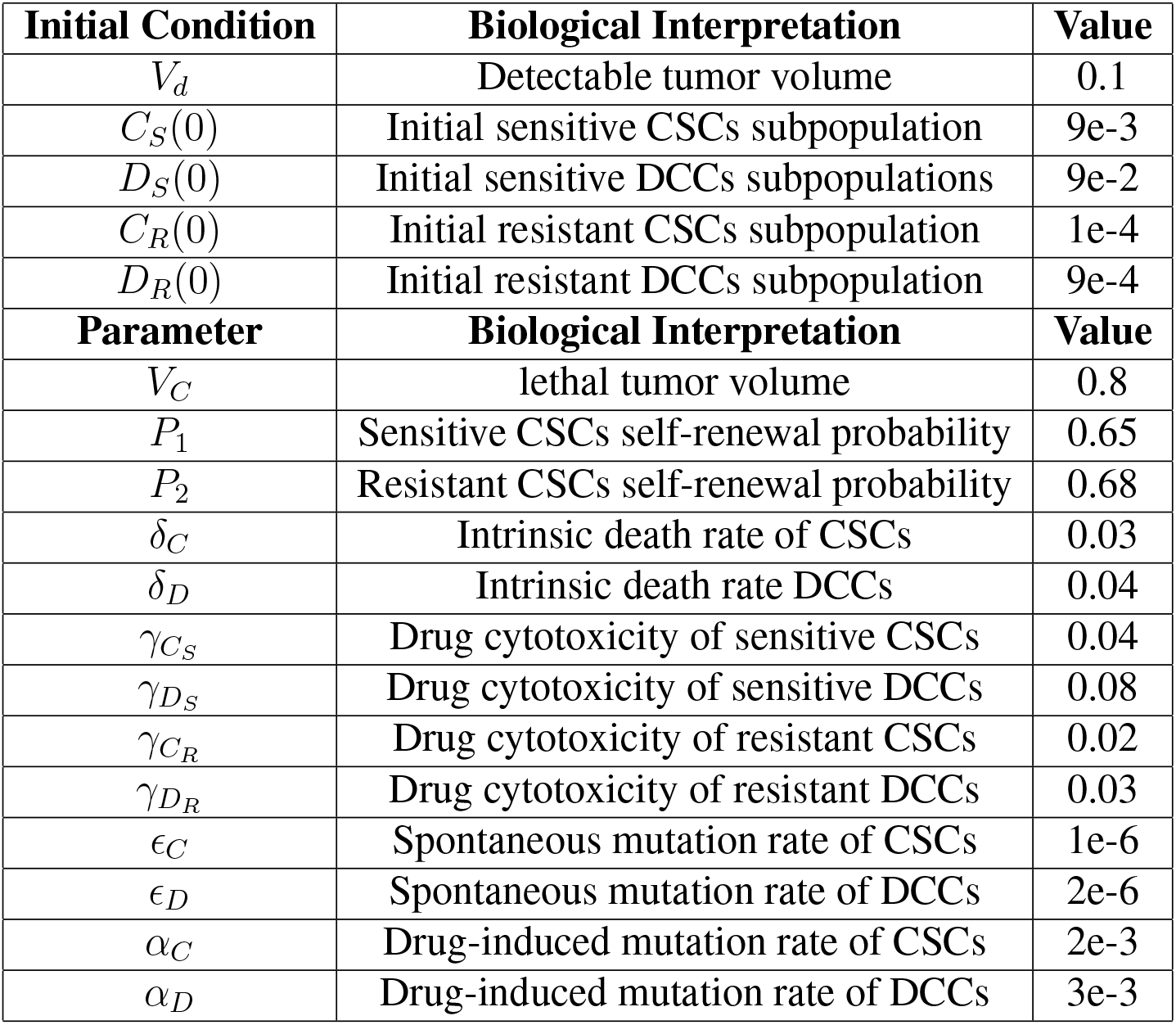
Initial condition and parameters for simulations in the Results section.

### Implementation Availability

All Matlab implementations are available at https://github.com/hyzkxo/Drug_Resistantce.git.

## Results

### Control Group

We first simulate the system under the no-treatment condition (effective dosage, *u*(*t*) = 0) to assess its behavior in untreated patients, confirming its alignment with common expectations. This case is denoted as control group in the rest of the paper. As shown in Figure 1, in the absence of drug administration, until the tumor volume reaches *V*_*C*_, the majority of the cell population consists of sensitive CSCs and DCCs, which implies spontaneous mutation alone can not result in drug resistance. The growth of tumor volume decelerates as it approaches the critical volume due to its limited carrying capacity. The enrichment of CSCs is spontaneous and continuous. At the end of the survival time of the untreated virtual patient, the proportion of CSCs is 2.3-fold greater than at the start of the simulation.

**Figure 1.**
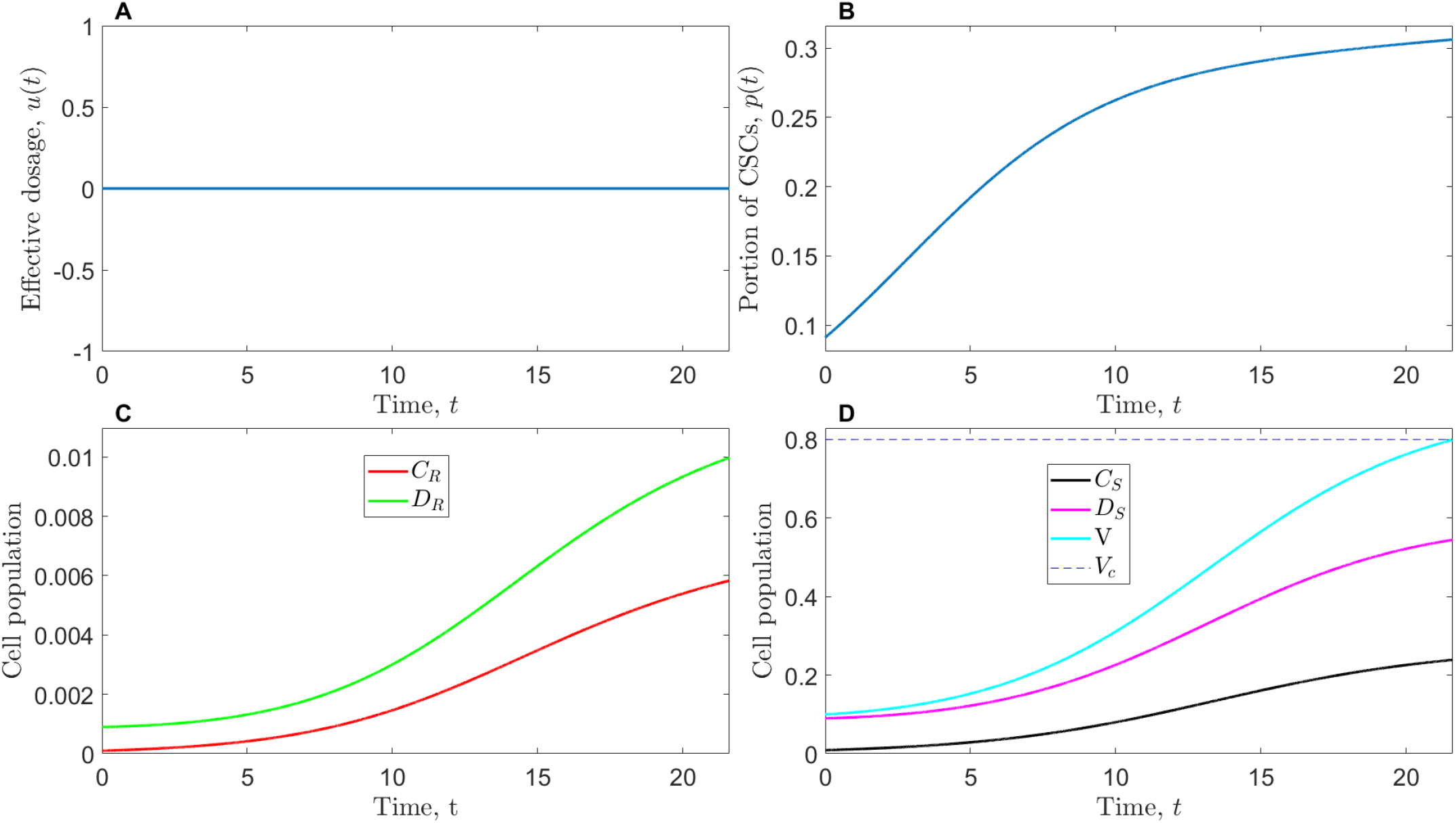
Control group, no-treatment condition, *u*(*t*) = 0. (A): Dosage function, *u*(*t*) = 0. (B): Proportion of CSCs in the whole cell population, *p*(*t*). (C): Resistant CSCs and DCCs subpopulations. (D): Sensitive CSCs and DCCs subpopulations and tumor volume, *V* (*t*).

### Constant Therapy

The first treatment simulated is constant therapy, where effective dosage is assumed to be constant value, *u*(*t*) = *u*_0_. Though this is the simplest form of therapy, it gives us some insights about therapy and drug resistance. As shown in Figure 2, compared with control group, survival time after constant therapy is 2.5-fold of that in control group. Consequently, we can verify the effectiveness of our virtual drug at a dosage of *O*(1), as it significantly prolongs the life of the virtual patient. CSCs enrichment is enhanced by constant therapy. Final proportion of CSCs is 1.3-fold relative to control group. The resistant cell (both resistant CSCs and DCCs) growth reaches 20-fold of that in control group. Unlike in control group, constant therapy results in strong cell population shift from sensitive to resistant phenotype. As shown in Figure 2.D, sensitive CSCs and DCCs gradually decays as tumor volume reaches lethal volume *V*_*C*_ = 0.8, while resistant CSCs and DCCs rapidly grow. The phenotype shift is mainly attributed to efficacy of drug on sensitive phenotype elimination and drug-induced transition from sensitive to resistant.

**Figure 2.**
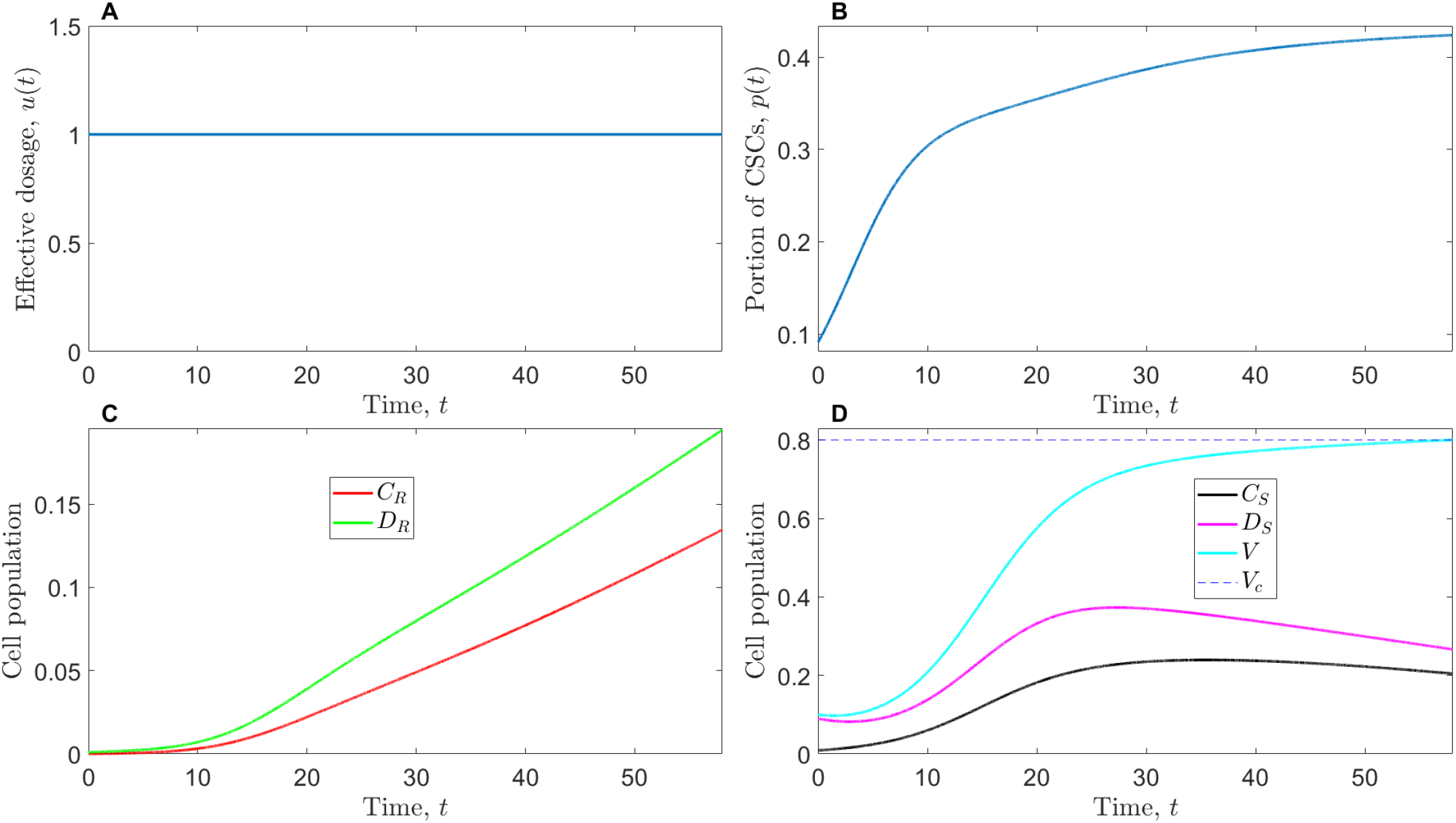
Constant therapy, *u*(*t*) = 1. (A): Dosage function, *u*(*t*) = 1. (B): Proportion of CSCs in the whole cell population, *p*(*t*). (C): Resistant CSCs and DCCs subpopulations. (D): Sensitive CSCs and DCCs subpopulations and tumor volume, *V* (*t*).

As shown in Figure 3, in constant therapy, the dosage, *u*_0_, positively regulates survival time, *t*_*C*_. Figure 3.A suggests that higher dosage triggers stronger suppression on tumor growth at early stage of treatment, but as resistance evolves in the tumor, treatment inevitably fails when drug response reduces and tumor volume reaches *V*_*C*_. Findings in Figure 3.B suggests that under constant therapy, survival time continues to increase with higher dosages. However, practical limitations arise from patients’ restricted ability to tolerate medication. In reality, indefinitely increasing dosages to extend patient survival time is not feasible due to physiological constraints.

**Figure 3.**
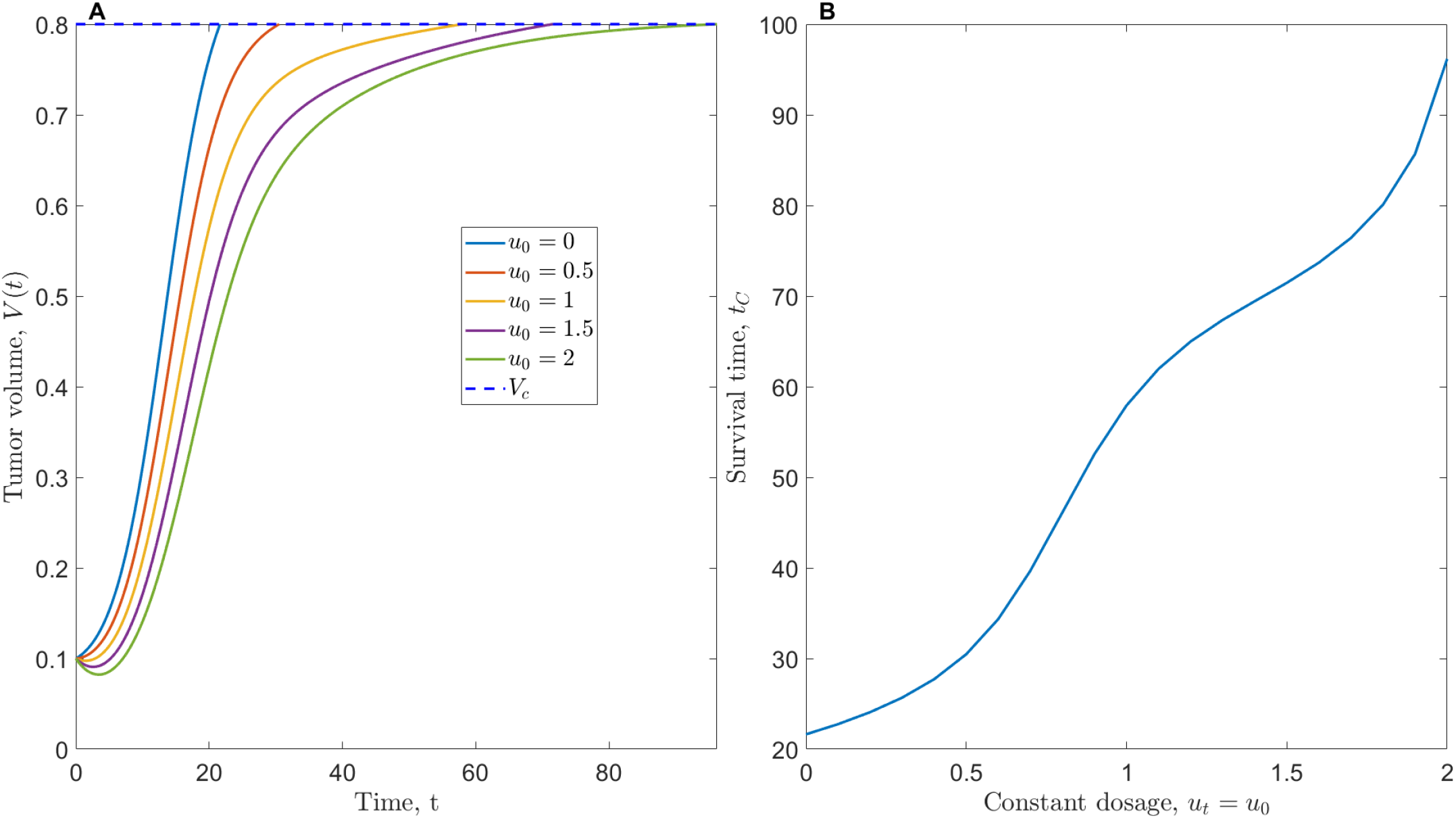
Effects of constant therapy dosage, *u*(*t*) = *u*_0_. (A): Tumor volume growth under different dosages. (B): Survival time, *t*_*C*_ under different dosages.

### Bolus Injection

Our virtual constant therapy faces two primary limitations: 1. In practice, maintaining an effective dosage in patients at a constant peak level is challenging due to factors such as metabolism [28]; 2. The rapid evolution of drug-induced resistance during constant therapy serves as the primary obstacle to its long-term efficacy. Therefore, we aim to develop an optimal treatment strategy with a more realistic dosage function and reduced drug-induced resistance in tumors. Furthermore, we expect that the new therapy will not impose additional cumulative dosage on virtual patients, because higher cumulative dosage reduces disease-free survival (DFS) rate [29].

Bolus injection refers to rapid administration of high dosage into the bloodstream. We aim to investigate the effectiveness of bolus injection as a treatment strategy. In our simulation, *u*(*t*) of bolus injection strategy is simplified as: 1. Injection is periodic with frequency, Ω, and conducted at the beginning of each cycle; 2. The effective dosage stays as constant level, *u*_*max*_, for a short period, Δ*t* and dosage is 0 for the rest of the time in each cycle; 3. To compare efficacy with constant therapy where constant dosage *u*(*t*) = 1 (Figure 2), we require *u*_*max*_ *·* Δ*t ·* Ω = 1 to obtain the same cumulative dosage in both constant therapy and bolus injection.

Simulation results with *u*_*max*_ = 12, Ω = 25, Δ*t* = 1*/*300 are summarized in Figure 4. Periodic bolus injection results in 17% increase in survival time compared with constant therapy at the same cumulative dosage level (Figure 4.D). The extension of survival time is primarily attributed to the significant suppression of sensitive CSCs and DCCs subpopulations by bolus injection, representing a 47% decrease compared to constant therapy, while corresponding resistant CSCs and DCCs subpopulations increase 66%. Bolus injection results in similar CSCs enrichment compared to constant therapy, with the CSCs proportion almost identical in both cases (final *p*(*t*) relative difference *<*3%). In summary, bolus injection extends the virtual patient’s survival time. The efficacy of bolus injection in extending the lifespan is attributed to its more effective elimination of sensitive phenotype CSCs and DCCs, which counteracts the enhanced drug-induced resistance associated with bolus injection protocols.

**Figure 4.**
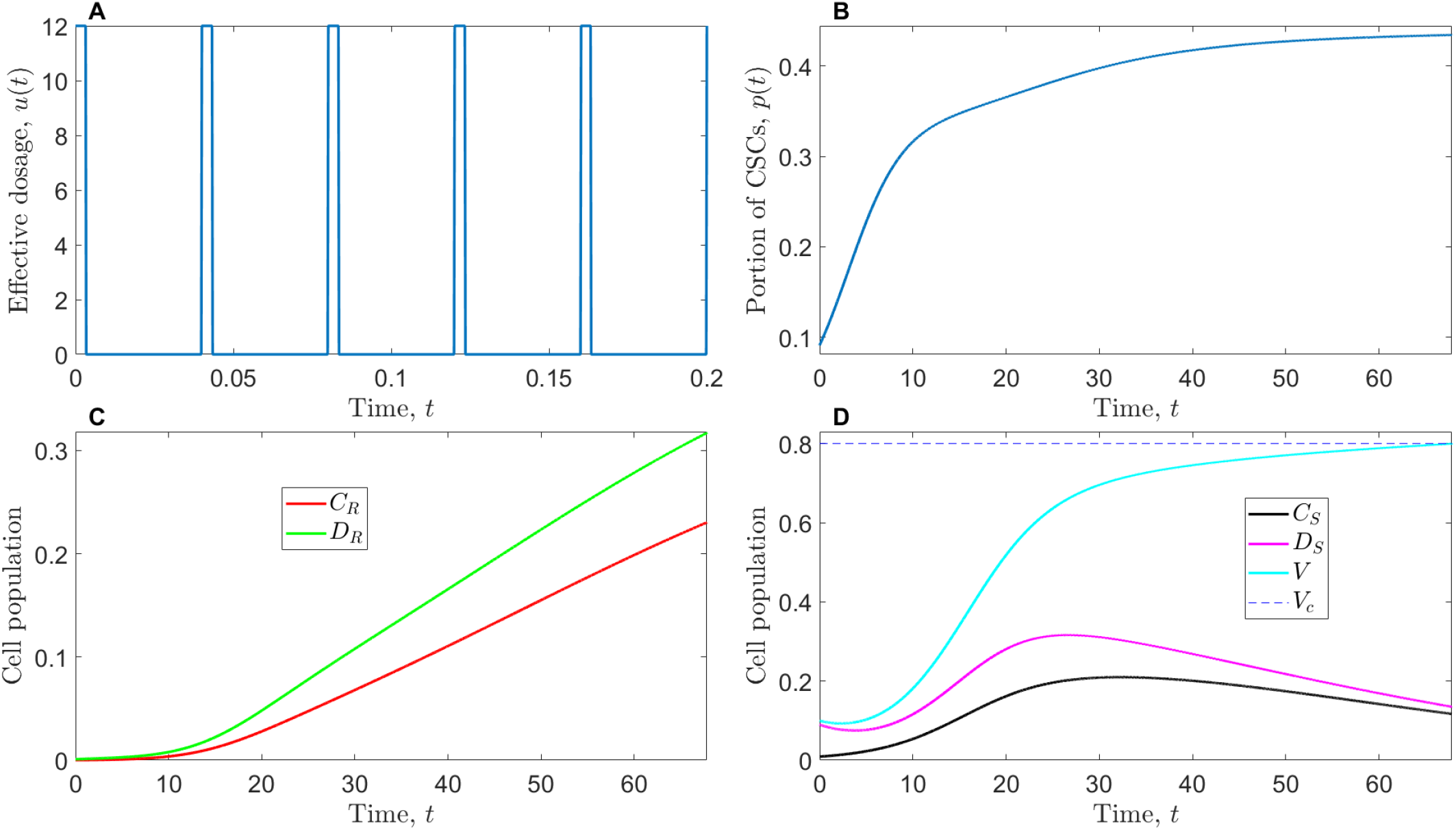
Periodic bolus injection. (A): Periodic dosage function, *u*(*t*) on interval [0,0.2]. Maximum dosage, *u*_*max*_ = 12; frequency of administration, Ω = 25; duration of maximum dosage, Δ*t* = 1*/*300. (B): Proportion of CSCs in the whole cell population, *p*(*t*). (C): Resistant CSCs and DCCs subpopulations. (D): Sensitive CSCs and DCCs subpopulations and tumor volume, *V* (*t*).

Similar to constant therapy, our next step is to understand the impact of maximum dosage in bolus injection while maintaining a fixed administration frequency. As demonstrated in Figure 5, at an administration frequency of Ω = 25, the escalation of maximum dosage does not consistently result in increased survival time; rather, it occasionally leads to a decrease in survival time. Figure 5.D suggests that the relationship between maximum dosage and survival time is bimodal with two peak survival times: Two peak survival times *t*_*C*_ = 66.6 and 68.5 are observed at *u*_*max*_ = 3.7 and 14.8 respectively, representing 9% and 12% relative increases compared to the maximum dosage of *u*_*max*_ = 12 in Figure 4. Treatment with higher maximum dosages yields slightly longer survival times compared to treatment with lower maximum dosages. This phenomenon may be explained through the interplay of maximum dosage and its duration: presuming a constant cumulative dosage at the same administration frequency, the duration of maximum dosage is proportional to the inverse of its magnitude. Consequently, near first peak *u*_*max*_ = 3.7 because the duration of dosage in each cycle is sufficiently long, the growth of sensitive phenotype is well suppressed, while near the second peak *u*_*max*_ = 14.8, though duration is short, high instant dosage also works well. Furthermore, a higher maximum dosage may exert stronger instant suppression on the growth of resistant phenotypes. Therefore, if the tumor volume approaches the lethal volume primarily due to resistant phenotypes, a higher maximum dosage is more effective and may marginally extend the patient’s life.

**Figure 5.**
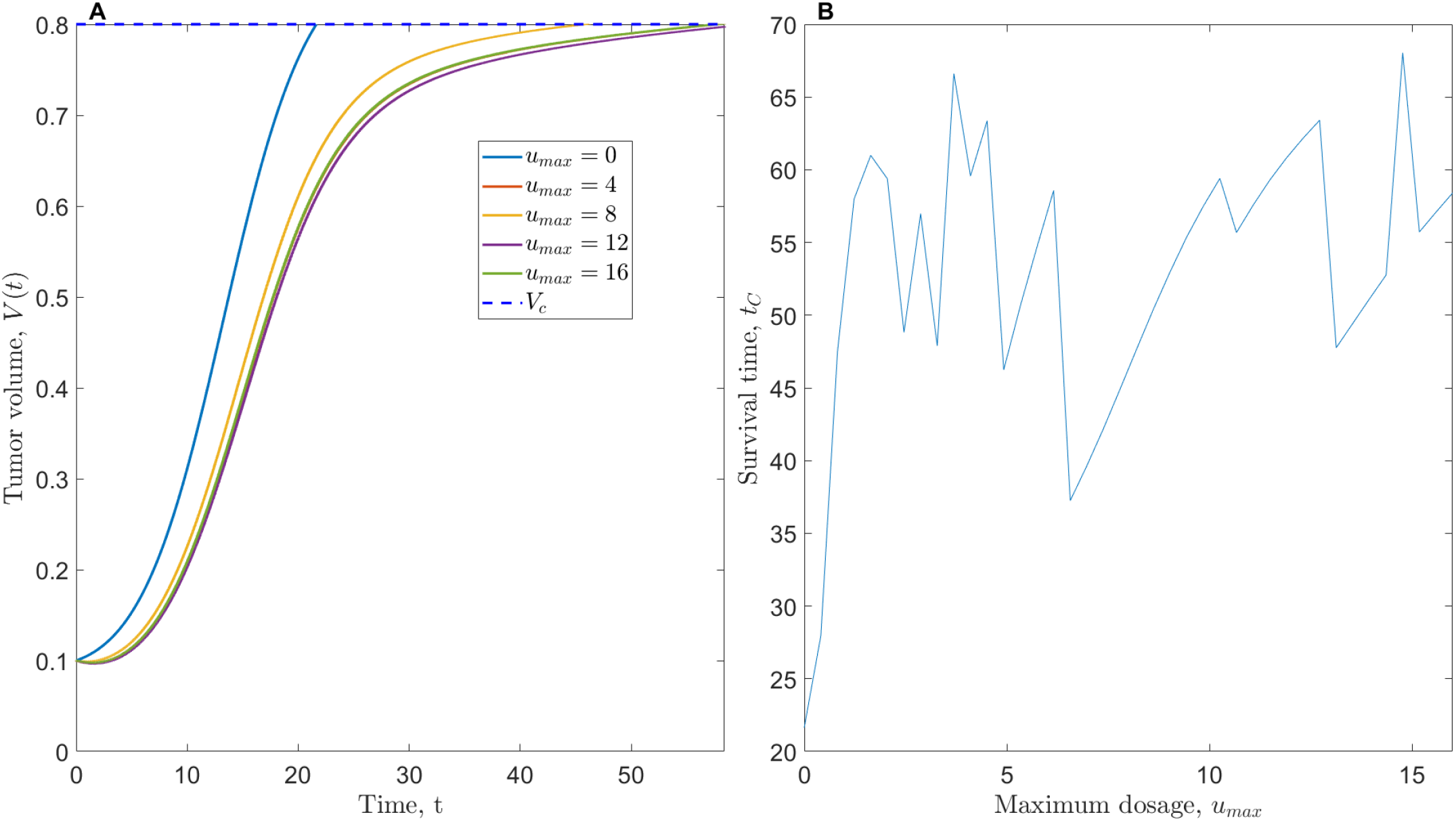
Effects of bolus injection maximum dosage, *u*_*max*_. (A): Tumor volume growth under different maximum dosages. (B): Survival time, *t*_*C*_ under different maximum dosages.

## Discussion

### Clinical Insights

Both simulations of constant therapy (Figure 2) and periodic bolus injection (Figure 4) suggest that chemotherapy promotes CSCs enrichment in the tumor. Compared with control group (Figure 1), as expected, constant therapy is particularly effective on sensitive DCCs compared with sensitive CSCs, which implies CSCs enrichment is a non-negligible component in the formation of drug-resistance in chemotherapy.

If the predicted bimodal relationship between bolus injection maximum dosage and survival time holds, two optimal treatment strategies emerge: 1. low maximum dosage with a long duration; 2. high maximum dosage with a short duration. Strategy 1 is more feasible as it demands less from the drug’s efficacy in achieving high effective dosages in patients. However, strategy 2 may result in a slight increase in survival time extension, which could benefit hospitals and patients with the capability to pursue it. Strategy 1 has an an illustrative application in breast cancer treatment, low-dose metronomic chemotherapy (LDMC): Krajnak et al. demonstrated that continuous administration of low-dose chemotherapy can be comparable in efficacy to traditional high-dosage chemotherapy, with the added advantage of enhanced inhibition of metastasis [30].

### Limitations and Future Work

Since all parameters in our simulations are dimensionless, it presents challenges in terms of validation against experimental data and accurately predicting the effects of various therapies. Thus, in our future work, it is important to seek actual data and incorporate parameters with appropriate units. Additionally, if we revisit original model Equation 2 the carrying capacity may be non-constant and significantly influenced by oxygen availability [31].

In our Model Formulation subsection, CSCs self-renewal probability, *P*_1_ and *P*_2_, remain constant. However, in certain types of solid tumors, it has been observed that the self-renewal rate is negatively regulated by the differentiated cell population [32], although the precise signaling pathway remains elusive. The introduction of such negative feedback could potentially alter the response of CSCs enrichment.

The dosage functions in constant therapy and bolus injection simulations lack realism due to the intricate relationship observed in practice between drug concentrations in plasma, tumor interstitium, and tumor cells [33]. The assumptions of constant effective dosage and instantaneous vanishing dosage in bolus injection are evidently oversimplified, serving the purpose of capturing essential behaviors under certain therapeutic conditions. In the follow-up work, more realistic effective dosage function should be utilized in the simulations.

In our model, cancer treatment and resistance evolution are based on a single type of drug. However, in clinical practice, chemotherapy often involves the use of multiple drugs in combination. For example, vincristine and prednisone is a standard combination to treat acute lymphoblastic leukemia [34]. Besides multi-drug resistance [35], drug-specific resistance exists. For instance, tamoxifen is widely utilized as the primary treatment for all stages of estrogen-receptor-positive breast cancer [36]. However, a significant portion, 40%, of patients die of tamoxifen resistance [37]. Even in cases of tamoxifen resistance, aromatase inhibitors are effective for breast cancer patients [38]. Therefore, the incorporation of multiple drugs into our model enables a more comprehensive and detailed quantitative discussion of drug resistance formation.

If we extend the simulation time interval, it is observed that the populations of sensitive CSCs and DCCs approach zero under multiple parameter sets and initial conditions. Does this imply that the system will asymptotically consist only of resistant CSCs and DCCs? In a framework model where CSCs and DCCs are not distinguished [15], Greene et al. mathematically proved that the sensitive phenotype will unconditionally vanish asymptotically, and the resistant phenotype is guaranteed to reach a maximum dimensionless population of 1. Can we similarly prove, under certain weak parameter constraints, that our proposed model Equation 4 exhibits analogous behavior?

### Bolus Injection

Besides LDMC, as mentioned earlier, various forms of bolus injection have been extensively utilized in breast cancer treatment. Speth et al. found weekly bolus injection of 4’-epi-adriamycin (E-ADM) maintain sufficient cellular drug concentrations in patients despite low plasma drug concentration [39].

While bolus injection has been validated as an effective chemotherapy method through both experimental studies and clinical practice across breast cancer and various other cancer types [40–42], it is limited in certain practical aspects. In breast cancer treatment, combined bolus injection of cyclophosphamide, methotrexate, fluorouracil incurs higher monthly expenses compared to infusion charges, primarily attributable to the elevated drug doses [43]. In the USA, 42% of patients declare bankruptcy due to medical expenses occurring after the second year following a cancer diagnosis [44]. Cost of therapy is a non-ignorant factor to select a chemotherapy method.

